# SPOTTER: Automated Tissue-Barcoding Platform for Spatial Proteomics and Phosphoproteomics

**DOI:** 10.64898/2026.06.08.730901

**Authors:** Yuanwei Xu, Hyeon-Cheol Park, Jason Li, Yuehan Liu, T. Mamie Lih, Cheng-Yu Lee, Xingde Li, Hui Zhang

## Abstract

Spatial proteomics aims to characterize proteome *in situ* with regional resolution, linking molecular states to tissue architecture and microenvironments. While recent advances have expanded our access to spatially resolved proteomics, major barriers in scalability and throughput remain. Building on our prior SPOT (Spatial Proteomics through On-site Tissue-protein-labeling) framework^1^, we introduce SPOTTER, a customizable robotic platform that enables automated, micron-scale spatial barcoding directly on intact tissue sections. SPOTTER barcodes proteins by labeling tissue proteins *in situ* to encode spatial origin across the whole tissue section. This strategy enables unbiased whole-tissue proteome mapping and, for the first time, spatial phosphoproteome profiling from intact sections. Coupled with high-resolution LC-MS/MS, SPOTTER achieves deep proteome and phosphoproteome coverage while preserving histological integrity. By replacing labor-intensive laser capture microdissection workflows and low-plex antibody arrays, SPOTTER provides a scalable route to high-plex spatial proteomics, resolving spatially distinct regions within a single experiment.

## Main

Proteins serve as the functional executors of almost all cellular activities. Protein modifications, spatiotemporal disparities, protein-protein interactions, proteolytic processing, and structural conformations collectively expand the diverse functional repertoire of gene expression. Despite extensive research and the generation of large-scale proteomic datasets from bulk tissue analyses, spatial proteomics data remain largely underrepresented. Bulk tissue proteomics often fails to capture the spatial heterogeneity of or cell-type-specific features of proteomes within the native tissue architecture, averaging signals across diverse cell types and masking critical cell-type-specific or regional variations in protein expression and function.

The limitations of bulk proteomics become particularly apparent when investigating dynamic regulatory processes within tissues. Many cellular activities are governed by spatially organized signaling mechanisms, where protein abundance, protein modifications, and molecular interactions vary across anatomical microenvironments. For example, the spatial distribution of phosphorylation events can reveal region-specific signaling networks that are critical for both physiological regulation and disease pathogenesis. The need for spatially resolved molecular measurements has been underscored by the rapid expansion of spatial transcriptomics technologies, including Visium^2^, MERFISH^3^, and Slide-seq^4^, which map RNA distributions across tissues at high resolution. However, protein abundance correlates only imperfectly with transcript levels and transcriptomic measurements^5^ cannot capture protein modification states, meaning that spatial transcriptomes cannot substitute for spatially resolved proteomic analyses. Despite this need, existing spatial proteomics techniques remain constrained by technical limitations.

Current spatial proteomics methodologies primarily fall into two categories: imaging-based techniques and bottom-up mass spectrometry (MS). Label-free mass spectrometry imaging (MSI) methods, such as matrix-assisted laser desorption/ionization (MALDI)^6–11^, secondary ion mass spectrometry (SIMS)^12,13^, and laser ablation electrospray ionization (LAESI)^14^,enable direct visualization of molecular distributions based on their mass-to-charge (m/z) ratios. Although these techniques offer the potential for high multiplexing without the need for pre-labeling, they often struggle with specificity and depth of coverage proteome or protein modifications. On the other hand, label-based multiplex imaging methods, including CODEX^15^, immunofluorescence (IF)^16^, imaging mass cytometry (IMC)^17,18^, and multiplexed ion beam imaging (MIBI)^19^, achieve high specificity by leveraging antibody or probe-based targeting of predefined antigens. However, these methods are inherently limited by antibody availability and their multiplexing capacity, which restricts the number of targets that can be simultaneously assessed.

Recent advancements in spatial proteomics, particularly through the integration of high-sensitivity mass spectrometers with techniques like laser microdissection (LMD), have pushed the envelope towards single-cell and even subcellular resolution. Approaches such as LMD-assisted NanoPOTS^20^ imaging and Deep Visual Proteomics (DVP)^21^ have demonstrated that it is possible to achieve detailed spatial maps of the proteome by combining precise tissue dissection with sensitive MS-based proteomics. Beyond LMD-based workflows, several non-LMD approaches have broadened the accessibility of spatial proteomics. Tissue micro-sampling with a 3D-printed scaffold (MASP)^22^ enabled micron-scale whole-tissue profiling, while Expansion proteomics (ProteomEx) achieved ∼160 µm lateral resolution via hydrogel-tissue transformation and micro-sampling without specialized equipment^23^. More recently, a microfluidics-based approach (PLATO) combined with transfer learning has more recently enabled high-resolution spatial proteomics of complex tissues^24^. Yet, these methods remain challenged by trade-offs among spatial resolution, proteome depth, and experimental throughput. While significant progress has been made in spatial proteomics, the field of spatial protein modifications has lagged behind. Phosphoproteomic analyses are particularly demanding due to the lower abundance and transient nature of phosphorylation events, compounded by the need for specialized enrichment protocols and the challenge of maintaining spatial information during sample preparation.

To bridge these gaps, we previously developed SPOT (Spatial Proteomics through On-site Tissue-Protein Labeling)^1^, a multiplexing method that integrates direct on-slide labeling of tissue proteins with bottom-up MS to achieve deep proteomic profiling while preserving spatial context. SPOT circumvented the laborious process of isolating specific tissue regions by enabling on-tissue, Tandem Mass Tag (TMT)-based direct labeling. This approach demonstrated the feasibility of whole-tissue mapping, highlighting spatial heterogeneity that is typically obscured in bulk analyses. Recognizing the need for high throughput, we have now advanced this approach by developing SPOTTER, an automated tissue-barcoding platform tailored for both spatial proteomics and phosphoproteomics. SPOTTER leverages a customizable robotic system that performs micron-scale spatial barcoding directly on intact tissue sections through the programmable deposition of TMT reagents. By encoding the spatial origins of proteins across an entire tissue section, SPOTTER not only enhances throughput but also facilitates the deep mapping of proteomic and phosphoproteomic landscapes without compromising histological integrity.

## Results

### SPOTTER printing workflow

SPOTTER integrates two functionally synchronized subsystems: a high-pressure nanoflow pump module for precision reagent delivery and a 3D-printing platform for spatially resolved deposition (Figure 1A). The printing module was engineered by retrofitting a commercial FDM printer with a fused silica capillary column as a microfluidic deposition nozzle, retaining the native XY motion control system for positional accuracy (Figure S1A, B). Custom G-code scripts coordinated reagent delivery with spatial positioning across predefined tissue regions. To minimize cross-contamination, the nozzle was flushed with 25% acetonitrile between deposition cycles. Each region was printed five consecutive times with 3-minute ambient drying intervals to ensure labelling efficiency. Following TMT labelling and quenching, labelled tissue was carefully lifted from the slide, lysed in 8M urea, and subjected to tryptic digestion.

**Figure 1.**
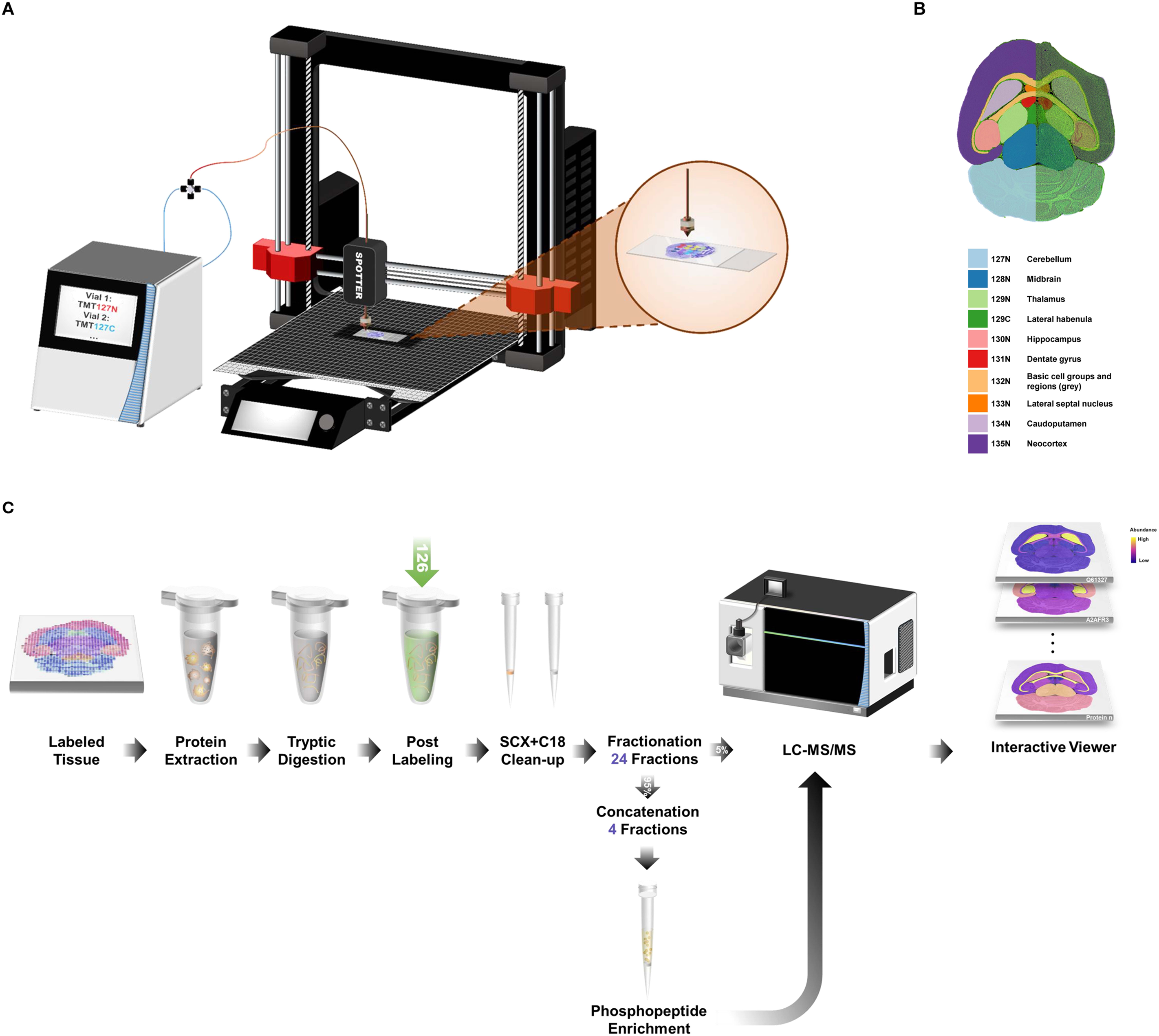
SPOTTER Proteomic Workflow. A) Set-up of SPOTTER for Direct Labeling of Tissue Proteins. SPOTTER integrates two functionally synchronized subsystems: a high-pressure nanoflow pump system for precision reagent delivery and a 3D printing platform for spatially resolved depositions, in which the thermoplastic extrusion assembly was replaced with a custom-made fused silica nano-flow liquid chromatography column as a microfluidic deposition nozzle. B) Murine brain atlas with region-to-tag assignments. Ten mouse brain regions were outlined and color-coded to illustrate their corresponding TMTpro-18plex tag assignments. C) SPOTTER workflow: (1) Annotation of brain regions on tissue sections. (2) Region-specific on-slide labeling using a custom printing pattern. (3) Tissue lift-off, lysis, and digestion. (4) Peptide-level TMT post-labeling to correct for unlabeled background. (4) Fractionation of labeled peptides. (5) Parallel LC-MS/MS analysis for global proteome and IMAC-enriched phosphoproteome. (6) Interactive web-based viewer for direct visualization of the proteomic distribution of proteins and phosphosites.

To achieve accurate, quantitative protein mapping on the whole-tissue level, the deposition of labeling reagents should follow a consistent fashion. SPOTTER’s printing performance was assessed by depositing a 10 × 10 grid pattern onto Polysine™ microscopic slides at an XY travel speed of 200 mm/s and a Z approach and retract speed of 16.67 mm/s, using a fluorescent solution (1% rhodamine B in 25% v/v acetonitrile) (Figure S1D). Nozzles with varying inner diameters (75 µm/ 30 µm/ 10 µm i.d.) and flow rates (1 µL/min, 0.2 µL/min, 0.1 µL/min) were tested to evaluate dot deposition uniformity and reproducibility. Post-printing, slides were imaged with a GenePix 4000B Microarray Scanner (Molecular Devices) to capture fluorescence signals. Acquired images were analyzed using Fiji/ImageJ (NIH) with the “Analyze Particles” tool to quantify morphological features of the printed dots, including their area, circularity, and diameter.

Our initial evaluation with a 75 µm-i.d. nozzle at 1 µL/min yielded printed dots with a median diameter of ∼ 859 µm and high reproducibility (coefficient of variation, CV = 2.58%). Switching to a 30 µm-i.d. nozzle at the same flow rate reduced the median dot size to ∼ 617 µm while maintaining low variability (CV = 1.69%). To further minimize the diameter of the printed dots, a 10 µm-i.d. nozzle was tested: at 0.2 µL/min, the median dot diameter decreased to ∼ 366 µm (CV = 0.91%), achieving a 41% reduction compared to the 30 µm nozzle. When the flow rate was further reduced to 0.1□µL/min (the lowest operating limit for Easy nanoLC pump) using the 10□µm nozzle, the median dot size decreased to ∼ 168□µm with a CV = 1.46%.

Alternatively, to further improve the resolution beyond the capabilities of the continuous-flow system, we developed a pin-based variant of the SPOTTER. In this configuration, reagent delivery is replaced by a solid pin with an inner capillary channel 15 µm i.d. (Figure S1C). The pin-based system makes direct contact with the substrate surface, enabling stable dispensing at sub-nanoliter volumes and reducing the minimum achievable dot diameter to ∼ 90 µm with a CV = 1.56%. Despite the wider inner channel relative to the 10 µm silica nozzle, the pin-based nozzle physically constrains lateral reagent spreading independent of droplet formation and surface wetting dynamics. This pin-based platform represents the next iteration of the system and is the subject of ongoing characterization.

### Spatial proteomics reveals region-specific protein programs

To evaluate the SPOTTER’s capacity for sub-tissue spatial proteomics in complex mammalian tissue, we analyzed four 7 µm-thick horizontal murine brain sections, leveraging the well-defined anatomical segmentation of the murine brain. Sections were imaged and annotated according to the Allen Brain Atlas^25^ to delineate ten anatomically defined regions: cerebellum (CB), midbrain (MB), thalamus (TH), lateral habenula (LH), hippocampus (HPF), dentate gyrus (DG), basic cell groups and regions (grey), lateral septal nucleus (LS), caudate putamen (CP), and cortex (CTX) (Figure 1B). Region-specific printing patterns were generated according to anatomical boundaries and implemented using a 30 µm inner-diameter nozzle (1 µL/min flow rate) to match regional scale. Following spatial labeling and quenching, entire sections were lysed in 8 M urea buffer and processed for downstream proteomic analysis.

To minimize interference from unlabeled peptides, an additional peptide-level labeling step was performed using an unused TMT channel distinct from those applied during on-slide spatial printing. Five percent of total peptides were analyzed for global proteomics, while the remaining fraction underwent IMAC enrichment for phosphoproteomic profiling (Figure 1C).

Across three sequential tissue sections, SPOTTER consistently quantified 8,025-8,398 proteins (24-fraction global proteomics), with pairwise inter-replicate Pearson correlations showed consistent protein profiling across the brain regions, using the cerebellum as a representative region (Figures 2A-B and Figure S2). These metrics establish that the spatial barcoding and sample preparation workflow introduces minimal replicate-to-replicate noise, a prerequisite for reliable downstream spatial inference.

**Figure 2.**
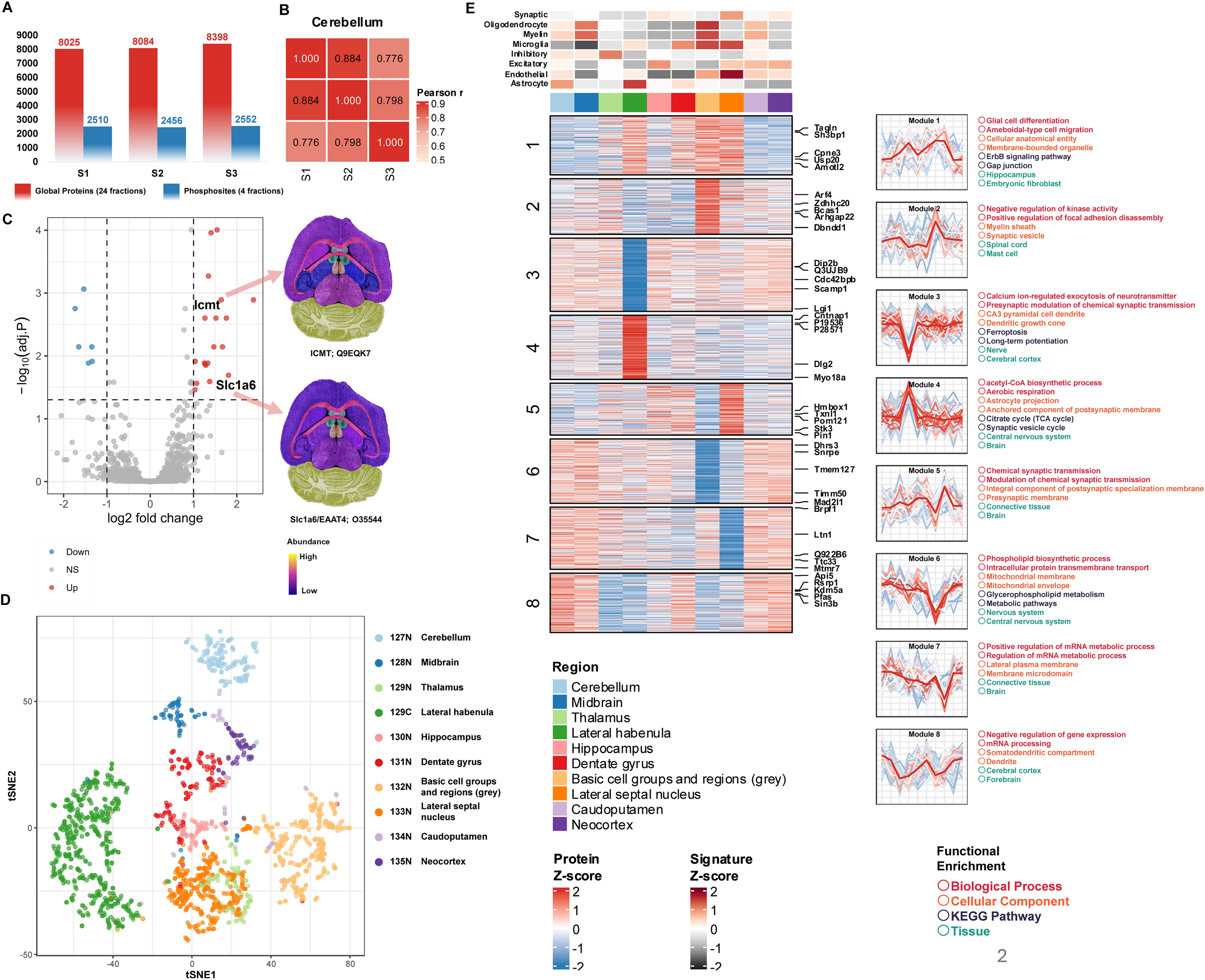
Spatial proteomics reveals region-specific protein programs. A) Number of proteins (global, 24 fractions; red) and phosphosites (4 fractions; blue) identified across three biological replicates (S1–S3) of murine brain sections. B) Pairwise Pearson correlation matrix of protein abundance values across the three cerebellum replicates. C) Differential abundance analysis for the cerebellum versus other brain regions. The volcano plot (left) shows significantly up- (red) and down-regulated (blue) proteins. Right panel displays region-resolved heatmaps for two example cerebellum-enriched proteins: Icmt (Q9EQK7) and Slc1a6/EAAT4 (O35544). E) Soft clustering of global proteomics data across ten brain regions using Mfuzz yields eight fuzzy clusters, displayed as Z-score heatmaps. The cell-type composition of each brain region is visualized through the expression levels of known cell-type marker proteins (Synaptic, Oligodendrocyte, Myelin, Microglia, Inhibitory, Excitatory, Endothelial, Astrocyte) as signature bars (top), illustrating region-specific cell-type identity. Cluster membership profiles are shown as line plots (center right) with representative proteins annotated (center left), and functional enrichment terms from biological pathway, cellular component, KEGG pathway, and tissue ontologies are annotated for each cluster (right).

To identify proteins enriched in individual brain regions, differential abundance analysis was performed using the limma linear modeling framework (each region versus the mean of all others; FDR < 0.05, |log2FC| > 1), with region-specific results visualized as volcano plots (Figure 2C and Figure S2). The cerebellum provided a stringent benchmark for regional specificity. Notably, ICMT (Protein-S-isoprenylcysteine O-methyltransferase; Q9EQK7), a protein with documented preferential expression in the adult cerebellum^26^, was among the most significantly enriched proteins, as was Slc1a6/EAAT4 (O35544), the glutamate transporter selectively expressed in Purkinje cells of the cerebellar cortex^27^. The spatial abundance heatmaps for these two proteins illustrate how region-enriched signals are distributed across the ten profiled areas, with cerebellar enrichment clearly resolved against a lower background in all other regions (Figure 2C). The recovery of these canonical, independently validated cerebellar markers and provided further support that SPOTTER accurately captures spatial biology rather than technical artifact.

To assess whether the full regional proteome structure captures anatomical identity, we performed t-SNE dimensionality reduction on the batch-corrected protein matrix. Proteins colored by brain region formed tight, region-specific clusters in the t-SNE embedding (Figure 2D), demonstrating that proteomic profiles are sufficiently distinct across regions to support unsupervised separation without any prior anatomical knowledge.

To identify higher-order spatial protein programs beyond individual region markers, we performed Mfuzz soft clustering on region-averaged, z-scored protein profiles, identifying eight co-variation modules with structured spatial enrichment patterns (Figure 2E). To link these proteomic modules to their cellular origins, we overlaid the expression levels of curated marker proteins spanning eight cell types (Synaptic, Oligodendrocyte, Myelin, Microglia, Inhibitory, Excitatory, Endothelial, Astrocyte)^28–32^, visualizing the cell-type composition of each region directly from the proteomic data. The resulting signatures aligned with established neuroanatomical cellular composition^25,28^, confirming that SPOTTER-derived modules recapitulate region-specific cellular architecture. The top five membership-weighted proteins per module were identified by ranking proteins on a combined score (Mfuzz membership × Pearson correlation to module mean profile), selecting those that most faithfully represent each module’s spatial trend.

### Spatial phosphoproteomics reveals decoupling of signaling from protein abundance across brain regions

Spatial phosphoproteomics is technically demanding on multiple fronts. Phosphopeptides are substoichiometric and typically require dedicated enrichment strategies, a substantially greater sample input is therefore required to achieve meaningful phosphosite coverage compared to standard proteomics. SPOTTER’s whole-tissue labeling strategy substantially increases the peptide input available for Fe^3+^-IMAC enrichment relative to microdissection-based workflows, enabling recovery of >2,500 phosphosites per experiment (Figure 1C, Figure 2A). More importantly, because global protein abundance is quantified in parallel from the same tissue section and the same TMT channels, SPOTTER provides the data needed to perform occupancy correction at scale across ten anatomical regions simultaneously.

For each phosphosite and each brain region, we computed a region-enrichment score defined as the difference between mean regional log2 abundance and the mean across all other regions (requiring ≥2 quantified samples within the region and ≥5 across other regions). Protein-level correction requires that the parent protein of each phosphosite be independently quantified in the global proteome, and that finite values are present in both matrices simultaneously within each sample. Of the 3,676 phosphosites identified across all three sections in the phosphosite matrix, 3,593 (97.7%) had a matched entry in the global proteome and were retained for protein-level correction. Because protein missingness compounds on top of phosphosite missingness, the joint completeness requirement substantially reduces the classifiable set: applying a ≥30% completeness threshold to the protein-corrected matrix (requiring finite values in ≥9 of 30 samples in both matrices) retained 548 sites, with the remaining 3,045 (84.7%) excluded at this step. These 548 protein-corrected sites were intersected with the 556 sites from the differential analysis, yielding the final set used for classification and visualization in the circular plot (Figure 3A) and the stacked bar chart (Figure 3B). Applying this framework classified a total of 223 significant phosphosites, including 181 regulation-dominant sites (red), whose spatial enrichment persists after protein-level correction, indicative of site-specific kinase activity independent of protein remodeling; 33 expression-dominant sites (orange), whose apparent regulation is mostly accounted for by changes in parent protein abundance; and 9 expression-masked sites (blue) exhibited phosphorylation changes that are only detectable following protein-level correction. The spatial distribution of these categories is visualized as a circular plot in which radial distance reflects the magnitude of differential phosphorylation (|log2FC|, capped at 4) and shaded sectors denote individual brain regions (Figure 3A).

**Figure 3.**
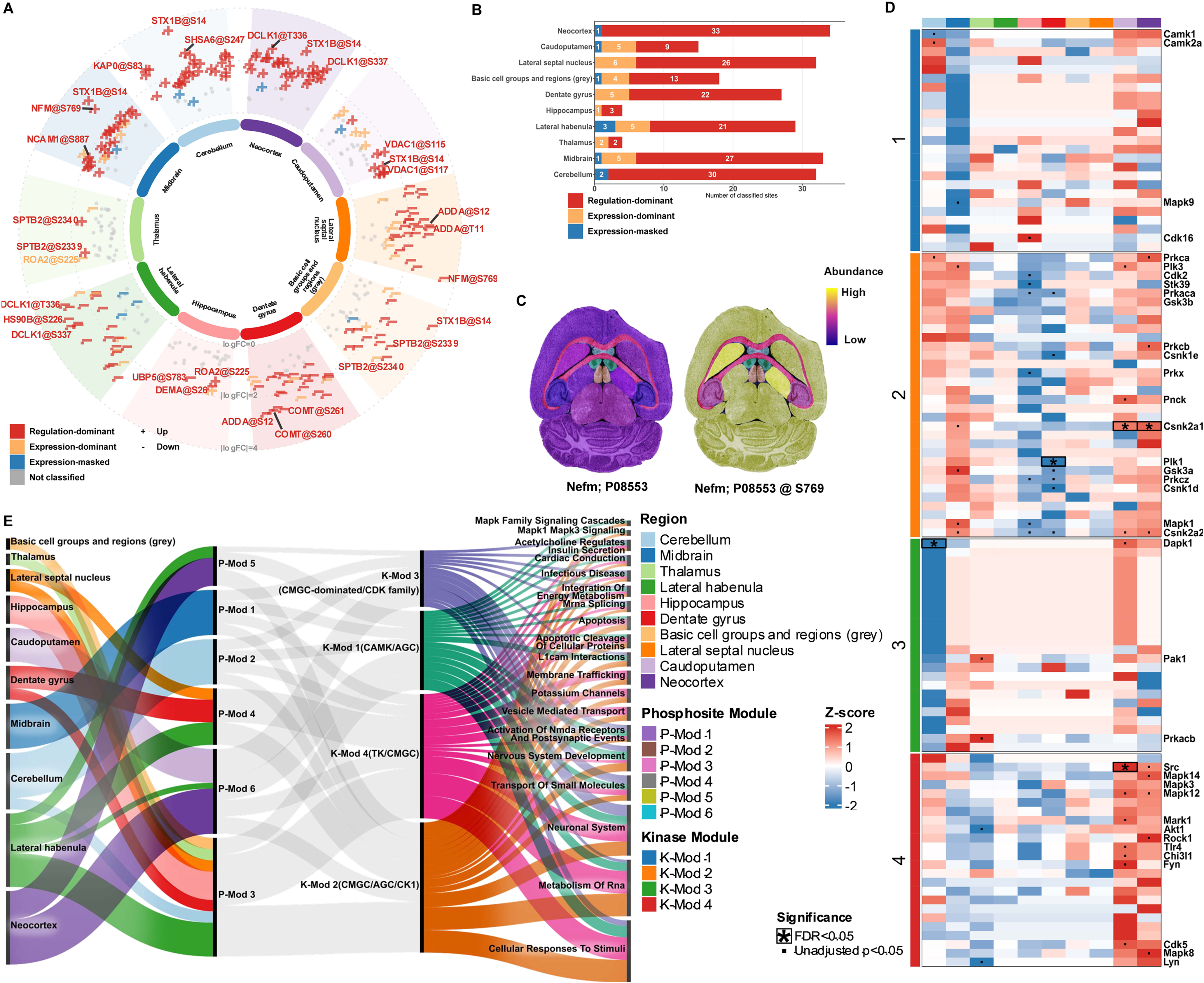
Decoupling phosphorylation signaling from protein expression. A) Spatial regulation of phosphosites across 10 mouse brain regions. Each point represents a phosphosite–region observation. Radial distance is proportional to |log₂ fold-change (FC)| (capped at 4). Protein-corrected sites are shown as + (up-regulated) or − (down-regulated) and colored by class: red = regulation-dominant, orange = expression-dominant, blue = expression-masked. Unclassified significant sites (adj. p < 0.05, |log2FC| ≥ 1) are shown as grey circles. The inner ring color denotes brain region. The top 3 sites per region are labelled. B) Stacked bar chart showing the number of phosphosites per brain region classified as regulation-dominant, expression-dominant and expression-masked. C) Example brain region heatmaps for Nefm (P08553) at the global protein level (left) and at a specific phosphosite (Nefm @ S769, right), demonstrating divergent spatial regulation between protein abundance and site-specific phosphorylation. D) Kinase activity enrichment analysis (KSEA) across 10 brain regions, summarized as four kinase modules (K-Mod 1–4) identified by k-means clustering of z-scored KSEA scores. Sites were required to be quantified in at least 4 regions; the top 50% most variable kinases were retained prior to clustering. Modules are displayed as z-score heatmaps with representative kinase families annotated. E) Sankey plot linking each brain region (left) to phospho-module (center-left) to kinase-module (center-right), and enriched reactome pathways (right), revealing module-specific signaling networks across the mouse brain.

An example of signaling-regulated phosphorylation is provided by Nefm (P08553). While the global protein abundance heatmap shows moderate enrichment in basic cell groups and regions (grey), the phosphosite at S769 displays a strikingly different spatial pattern, with preferential enrichment in the cortex and cerebellum (Figure 3C), demonstrating that site-specific phosphorylation can encode spatial information that is entirely invisible in total protein abundance maps.

To infer the upstream kinase programs responsible for spatially regulated phosphorylation, we applied Kinase Substrate Enrichment Analysis (KSEA) to region-enrichment scores using OmniPath enzyme-substrate relationships for Mus musculus. Region-wise kinase co-activity networks are shown for all ten brain regions (Figure S4). The kinase profiles (kinases quantified in ≥4 regions) were clustered into four modules by Mfuzz soft clustering of z-scored KSEA scores (Figure 3D). K-Mod 1 (CAMK/AGC, led by CaMK2a) is enriched in hippocampus and cortex, consistent with roles in synaptic plasticity^33^. K-Mod 2 (CMGC/AGC/CK1, including Ck2 and Plk1) reflects transcriptional and cell-cycle regulatory activity^34^. K-Mod 3 (CMGC; Mapk1, Dapk1) is enriched in midbrain and basic cell groups and regions (grey). K-Mod 4 (TK/CMGC; Mapk14, Mark1, Dyrk1a, Cdk8) concentrates in the thalamus and lateral habenula, consistent with circadian and tau regulation^35^. A Sankey diagram integrating kinase module membership (Figure 3D), phospho-module membership (Figure S3), enriched Reactome pathways, and brain regions provide a systems-level overview of spatial signaling architecture (Figure 3E). Each kinase module showed a distinct connectivity profile. K-Mod 1 (CAMK/AGC) flows primarily into P-Mod 2 (cerebellum; neuronal system, RNA metabolism, mRNA splicing) and P-Mod 6 (neocortex/caudoputamen; neuronal system, NMDA receptor activation), with additional flow into P-Mod 5 via Mapk1/Mapk3 signalling and L1cam interactions. K-Mod 2 (CMGC/AGC/CK1) is the most broadly distributed, connecting to P-Mod 1 (midbrain), P-Mod 3 (cerebellum; cellular stress and apoptosis), and uniquely to P-Mod 4 (dentate gyrus, lateral habenula; membrane trafficking, potassium channels, vesicle transport). K-Mod 3 (CMGC/CDK) carries the largest flow into P-Mod 6 and the strongest contribution to P-Mod 1 among all modules. K-Mod 4 (TK/CMGC; Src, Fyn, Egfr, Ptk2) mirrors K-Mod 2 in connecting to P-Mod 4 but through a distinct receptor tyrosine kinase composition. Together, these results demonstrate that spatial phosphorylation in the brain is organized into region-specific signaling programs with distinct directional patterns that reflect the functional specialization of each anatomical region. These findings further establish that SPOTTER provides sensitivity, spatial resolution, and analytical framework required to decode the signaling layer of tissue organization.

### An interactive browser enables real-time exploration of spatial proteomics and phosphoproteomics data

Deep spatial proteomics datasets present an interpretive challenge that scales with their coverage: with >8,000 proteins and >2,500 phosphosites mapped across ten brain regions (Figure 1C and 2A), meaningful biological interrogation requires tools that allow researchers to move fluidly between the global patterning of the dataset and the spatial behavior of individual proteins or modification sites. Static heatmaps capture summary views but cannot support the hypothesis-driven, protein-by-protein exploration that translates a large dataset into biological insight. To address this, we developed a GitHub-hosted interactive mouse brain protein and phosphosite heatmap viewer that makes the full SPOTTER dataset publicly accessible in real time (Figure 4).

**Figure 4.**
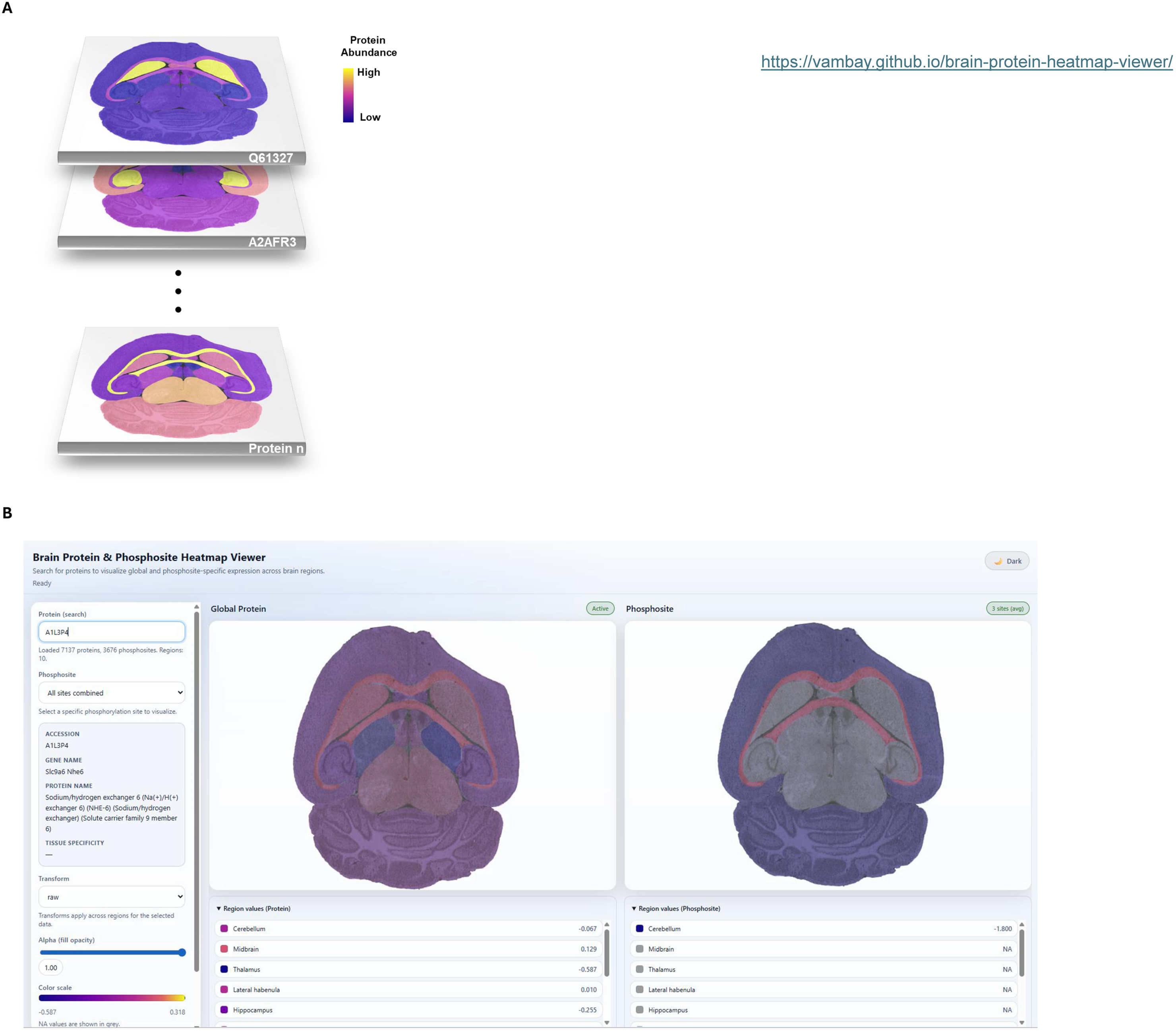
Interactive brain protein and phosphosite heatmap viewer. A) Protein abundance values derived from global TMT-based spatial proteomics are mapped onto annotated mouse brain regions and rendered as color-scaled heatmaps over a reference atlas, shown here as stacked layers across all quantified proteins. B) A GitHub-hosted interactive viewer enables real-time exploration of region-resolved protein and phosphosite expression. Users can search by UniProt accession or gene name, select individual phosphorylation sites or combine all sites, apply data transforms, and adjust display parameters including opacity and color scale. Quantified values for key brain regions are displayed as ranked tables alongside each heatmap. Missing values are shown in gray.

The viewer renders region-resolved protein and phosphosite abundance as color-scaled heatmaps overlaid on a mouse brain reference atlas at the horizontal pane, providing immediate spatial context for any queried protein. Users can search by UniProt accession or gene name to retrieve the global protein abundance map alongside the phosphosite map for the same protein, displayed side by side for direct comparison of protein-level and modification-level spatial patterns (Figures 4A-B). For phosphosite queries, individual modification sites can be selected or combined across all detected sites, enabling inspection of both site-specific and aggregate phosphorylation signatures. Display parameters including opacity and color scale are adjustable in real time, and region-specific quantification values are listed alongside each heatmap. Regions with missing quantification are displayed in grey, preserving transparency about data completeness across the ten profiled brain regions.

The interpretive value of juxtaposing protein and phosphosite maps in the same interface is illustrated directly by proteins such as Nefm (P08553): the global abundance heatmap and the S769 phosphosite map display strikingly divergent spatial patterns (Figure 3C), a discordance that would be invisible from either view alone and that motivates the protein-level correction framework described above. More broadly, the full value of the data is realized through open, interactive access that allows the broader community to query the resource against their own biological questions. The viewer, alongside median-normalized and log2 transferred raw data and analysis code, is publicly available at https://github.com/VamBay/brain-protein-heatmap-viewer.

## Discussions

Our study introduces SPOTTER, an automated tissue-barcoding platform that advances spatial proteomics and phosphoproteomics through programmable, micron-scale deposition of TMT reagents directly onto intact tissue sections. This approach overcomes key limitations of existing spatial proteomic methods, including scalability and throughput, while avoiding the biases inherent to antibody-based or LMD-based strategies: antibody-based strategies are limited to predefined targets, constrained by antibody availability, and incompatible with omics-scale discovery; while LMD-based approaches are sequential, require prior structural guidance, and are inherently limited to discrete dissected regions rather than whole-tissue landscapes. In terms of throughput, SPOTTER enables simultaneous barcoding of up to 18 spatially distinct regions in one experiment through TMTpro-18plex multiplexing, while LMD-based approaches require sequential laser cutting and sample collection. In terms of scalability, the programmable deposition architecture allows rapid reconfiguration of spatial sampling patterns without hardware modifications, making it readily adaptable to tissues of varying size and complexity. With respect to resolution, the deposition spot diameter of SPOTTER (∼168 µm with the flow-based configuration) is comparable to the spatial resolution typically employed in LMD-based proteomics workflows (100-200 µm). The pin-based variant (∼90 µm) further approaches the higher-resolution LMD range (∼20-50 µm. SPOTTER thereby enables unbiased, whole-tissue mapping of the proteome and phosphoproteome, achieving deep coverage of >8,000 proteins and >2,500 phosphosites per experiment. This substantially exceeds numbers of proteins typically reported for region-scale microdissection workflows^21,36^, while enabling spatial phosphoproteomic measurements that remain largely inaccessible to existing spatial proteomics approaches.

The application of SPOTTER to murine brain sections revealed distinct regional proteomic signatures and phosphorylation gradients. The phosphoproteomic analysis demonstrates that spatial phosphorylation is not simply a scaled reflection of protein abundance variation. Protein-level correction across ten brain regions revealed that a substantial fraction of phosphosites is classified as kinase-regulated: their spatial patterns diverge from those of the parent protein, indicating active kinase regulation rather than passive co-variation with protein levels. The kinase activity patterns inferred by KSEA show strong concordance with established functional neuroscience, providing orthogonal biological validation of the analytical framework.

While SPOTTER represents a significant technological leap, several avenues remain for further optimization. Future efforts could focus on refining deposition parameters to reduce variability at ultra-low flow rates and exploring the integration of additional multiplexing channels to enhance spatial resolution even further. The spatial resolution of SPOTTER is defined by the deposition spot diameter, which in the present study is optimized for anatomically defined brain regions. Spot dimensions are a tunable parameter of the deposition architecture rather than a fundamental constraint. Beyond spot dimension, effective spatial resolution is also influenced by inter-dot spacing and lateral diffusion of labeling reagents across the tissue surface. In the present implementation, lateral diffusion is mitigated using TMT reagents formulated in a high-acetonitrile solution, which under ambient deposition conditions evaporate rapidly and restrict lateral spread. Adaptation of SPOTTER to alternative tagging chemistries with slower evaporation kinetics would require careful characterization of reagent spreading behavior. Future engineering iterations can access finer spatial scales as advances in low-input sample preparation and mass spectrometry sensitivity expand the lower bound of detectable peptide input per spatial unit. Additionally, expanding the application of SPOTTER to other tissue types and disease models will be essential for fully realizing its potential in both basic research and clinical diagnostics. The present study uses fresh-frozen murine brain tissue as the primary analytical substrate. This choice reflects the high molecular integrity achievable with fresh-frozen material, the availability of well-defined anatomical reference atlases for murine brain, and the extensive prior proteomic, transcriptomic, and functional characterization of this tissue, which collectively make it an ideal system for method development and validation. Brain regions are unambiguously delineated anatomically and are known to differ substantially in their proteomic and signaling states, providing a stringent benchmarking ground for a spatial omics platform. The compatibility of SPOTTER with formalin-fixed paraffin-embedded (FFPE) tissue, as demonstrated *a priori* in (SPOT^1^), substantially broadens its translational potential given that FFPE is the standard archival format in clinical pathology and that vast repositories of clinically annotated FFPE material exist. Rigorous characterization of SPOTTER performance on FFPE tissue across a range of fixation durations and archival ages will be an important prerequisite for deployment in clinical research settings.

In summary, SPOTTER fills a distinct gap in the spatial omics toolkit, delivering unbiased, quantitative, site-specific proteomics and phosphoproteomics across anatomically defined spatial positions at a proteome depth unattainable by existing spatial methods. Its application to the murine brain reveals biologically coherent, spatially restricted kinase activity patterns that are inaccessible to both bulk proteomics and protein-level spatial analyses, establishing the platform as a foundational approach for the quantitative interrogation of signaling organizations in biological and clinical contexts, opening new horizons for the extensive understanding of molecular programs, cellular communication, plasticity, and regulatory dynamics.

## Materials and Methods

### SPOTTER Instrument Design and Fabrication

SPOTTER was developed in two operational configurations to accommodate different throughput and resolution requirements. The first configuration integrates a high-pressure nanoflow pump system for precision reagent delivery and a 3D printing platform for spatially resolved deposition. The pump module employs an Easy nLC 1200 (Thermo Fisher Scientific), which combines a 48-vial autosampler with temperature-controlled (4°C) storage, dual pressure-resistant pumps (1,200 bar maximum operating pressure; 100-1,000 nL/min flow range), and programmable gradient control. The printing module was engineered by retrofitting an Original PRUSA MK4 3D printer, wherein the thermoplastic extrusion assembly was replaced with a liquid-handling system. Critical modifications included disabling the print head’s heating element and embedding a fused silica nano-flow liquid chromatography column (75/30/10 µm inner diameter, 100 mm length; CoAnn Technologies) as a microfluidic deposition nozzle. The printer’s native motion control system features precise XY stepper motors (0.9° step angle, 1/16 microstepping) and polished steel rod linear bearings, enabling precise tip positioning during reagent deposition with a theoretical positional resolution of 5 µm. The two subsystems were bridged with a stainless-steel mixing tee (150 µm inner diameter; Thermo Fisher Scientific). In the second configuration, the fused silica nozzle was replaced with a ceramic capillary pin (OD 0.08 mm / ID 0.015 mm, LabNEXT), which enables direct contact-based reagent transfer onto the tissue surface and eliminating constraints imposed by pump flow limits and fluidic dead volume delays.

### Tissue Sample Collection and Preparation

Fresh-frozen horizontal mouse brain sections (7 μm thickness) were obtained from Zyagen (San Diego, California; Catalog # MF-201-HS). Prior to staining, sections were equilibrated to room temperature (20-22°C) and rehydrated by submerging slides in 1X phosphate-buffered saline (PBS; Thermo Fisher Scientific, 10010023) for 5 minutes to remove residual embedding medium and restore tissue hydration. Excess PBS was carefully removed using a low-pressure nitrogen stream to avoid tissue detachment.

To visualize cellular nuclei and anatomical landmarks, sections were stained with 0.1% Mayer’s hematoxylin (Sigma-Aldrich, Catalog # MHS32), a regressive stain ideal for differentiating nuclear chromatin. Hematoxylin was applied to fully cover the tissue section (500 μL/slide) and incubated for 5 minutes at room temperature. Unbound stain was removed by rinsing slides under running tap water gently for 2 minutes, removing excess dye while preserving tissue integrity. To enhance nuclear contrast and stabilize the hematoxylin stain, slides were immersed in 0.005% ammonium hydroxide (NH_4_OH) solution (prepared in 50 mL conical tubes) for 1 minute. This step neutralizes acidic residues, converting the hematoxylin’s reddish hue to a permanent blue-black signal. Following bluing, slides were rinsed twice in distilled water to eliminate residual NH_4_OH and prevent over-alkalization. Stained sections were gently dried under a controlled nitrogen stream (5-10 psi) to minimize oxidation or crystallization artifacts. To preserve spatial proteomic compatibility, slides were stored at -80°C in airtight slide boxes until further use. No coverslip was applied to avoid physical compression of the tissue or interference with downstream SPOTTER workflow.

### Tissue Sample Annotation and Pattern Gcode Conversion

The hematoxylin-stained mouse brain section was first imaged at 40X magnification using bright-field microscopy to capture baseline histological morphology. Prior to reference dots deposition, the slide was scanned on a GenePix 4000B microarray scanner (Molecular Devices) at two wavelengths (532 nm and 635 nm) with a resolution of 10 µm/pixel to establish pre-deposition spatial compositions. A grid of 19 × 48 reference markers (912 total dots) was robotically deposited onto the tissue using a solution containing 0.1% Mayer’s hematoxylin, acetonitrile, and ddH2O (2:1:1 v/v) to create spatial reference points. Post-deposition, the slide was re-imaged under identical GenePix 4000B settings to register the reference dots relative to tissue anatomy. Tissue architecture was annotated using data from the Allen Mouse Brain Atlas and pre-existing MRI images to align anatomical regions with reference dots. Coordinates corresponding to atlas-matched regions of interest (ROIs) were computationally extracted and converted into individual Gcode files using custom scripts. These Gcodes defined the spatial coordinates and printing parameters for subsequent TMT depositions.

### Tissue Sample Labeling Using TMT

Each TMT reagent (Thermo Scientific) vial was carefully opened, and the contents were gently suspended using the recommended volume of anhydrous acetonitrile by the manufacturer. The reagent was mixed thoroughly to ensure complete dissolution.

Suspended TMT reagents were reconstituted (TMT final concentration was 10µg/µL in 100mM HEPES, 25 % v/v acetonitrile) and printed directly to areas of interest using SPOTTER. The same procedure would be repeated for a total of 5 times, and between each pipetting, sections with labeling reagent would be left to air-dry. After labeling the sections of interest 5 times, 5% hydroxylamine was applied in a similar fashion to the whole tissue to quench the labeling.

### Tissue Lysis, Digestion and Post-digestion Labeling

Labeled tissue samples were scraped off the slides using a scalpel and transferred to microcentrifuge tubes. Samples were subjected to 8 M urea lysis buffer (8 M urea, 75 mM NaCl, 50 mM Tris-HCl, pH8). Enzymatic tryptic digestion was performed as previously described^37^. Digested tissue samples were cleaned up by SCX tip, desalted by C18 StageTip, and dried using Speed-Vac. For post-digestion labeling, dried peptides were resuspended in 100 mM HEPES and peptide concentration was measured by NanoDrop (Thermo Scientific). Peptides were then reacted with TMTpro 126 reagent at a 1:1.5 (reagent:peptide, w/w) ratio with shaking for 3 h at room temperature to cap remaining unlabeled primary amines.

### Basic Reverse-Phase Fractionation

TMT-labeled peptides were fractionated by high-pH reversed-phase chromatography on a ZORBAX Extend-C18 column (2.1 × 100 mm, 1.8 μm; Agilent) using an Agilent 1260 Infinity HPLC system. Mobile phase A was 10 mM ammonium formate (pH 10) and mobile phase B was 10 mM ammonium formate in 90% (v/v) acetonitrile (pH 10). Peptides were separated at 0.2 mL/min using the following gradient: 2% B for 10 min; 2-8% B over 5 min; 8-35% B over 85 min; 35-95% B over 5 min; and 95% B for 25 min. Eluate was collected into a 96-well plate and concatenated into 24 fractions. For global proteome analysis, 5% of each fraction was aliquoted, dried in a vacuum concentrator (SpeedVac), and stored at -80 °C. The remaining 95% was further concatenated into four fractions, dried in a SpeedVac, and stored at -80 °C for downstream PTM enrichment and LC-MS/MS analysis.

### IMAC Enrichment for Phosphopeptides

Dried peptides were suspended in 3%(v/v) ACN, 0.1%(v/v) TFA and peptide concentration was measured using Nanodrop (Thermo Scientific). Aliquots of ∼10 µg peptides were then reconstituted into 20 µL of 80%(v/v) ACN, 0.1%(v/v) TFA (peptide concentration controlled at ∼0.5 µg/µL) for the following IMAC enrichment.

IMAC procedure was performed using Fe^3+^-NTA agarose beads that were freshly prepared using Ni^2+^-NTA agarose beads (QIAGEN, cat no. 30210) as previously described^38^. Samples constituted in 80%(v/v) ACN, 0.1%(v/v) TFA were incubated with 60 μL of 5% (v/v) Fe^3+^-NTA agarose beads to conjugate for 30 min at RT. After conjugation, the supernatant containing unbounded peptides was collected by centrifugation. The beads with conjugated peptides were carefully transferred onto C18 Stage Tip and washed three times of 200 µL 80% (v/v) ACN, 0.1% (v/v) TFA. The washes were combined with supernatant for future use. The peptides conjugated to beads were eluted with 80 μL potassium phosphate buffer (500 mM KH2PO4, pH7) for 3 times and 50% (v/v) ACN, 0.1% (v/v) FA. Eluted phosphopeptides and unbounded peptides from IMAC were dried and stored at -80 °C for LC-MS/MS analysis.

### LC-MS/MS Analysis

Proteomic analysis was performed using tandem mass tag (TMT) labeling coupled with data-dependent acquisition (DDA) on an Orbitrap Ascend mass spectrometer (Thermo Fisher Scientific) interfaced with an Evosep One liquid chromatography system (Evosep). Peptide separation was achieved using a 15 cm × 150 µm inner diameter PepSep C18 reversed-phase column (Bruker, Cat. #1893474) packed with 1.5 µm particles. Chromatographic separation employed the Evosep One standardized 15 samples-per-day (SPD) method, utilizing an 88-minute gradient. The mobile phase consisted of solvent A (0.1% formic acid in water) and solvent B (0.1% formic acid in acetonitrile).

Electrospray ionization was performed at 1.9 kV using a 150 µm outer diameter × 30 µm inner diameter stainless steel emitter connected to the analytical column, with the ion transfer tube temperature maintained at 300°C to optimize ion desolvation. Full-scan MS1 precursor spectra were acquired in profile mode on the Orbitrap Ascend at a resolution of 120,000, covering a mass range of 400–1400 m/z with an automatic gain control (AGC) target of 1.2 × 10^6^ and an automatically optimized maximum injection time over the 88-minute gradient. Data-dependent acquisition triggered HCD-MS/MS spectra (collision energy: 35%; isolation window: 0.7 m/z) for the most intense precursors, with MS2 spectra collected in centroid mode at 30,000 resolution (AGC: 5 × 10^4^ max injection time: 59 ms) within a 3-second duty cycle. Monoisotopic precursor selection (peptide mode) and charge state filtering excluded unassigned 1+, 7+, 8+, and >8+ ions, while dynamic exclusion (45 s; ±10 ppm mass tolerance) minimized redundant fragmentation of previously analyzed ions.

### Database Search and Data Preprocessing

Raw files were processed in Proteome Discoverer 3.0 (Thermo Fisher Scientific) using SEQUEST HT to search against the Mus musculus UniProt proteome (downloaded December 19, 2023), incorporating both standard trypsin cleavage (C-terminal to lysine/arginine, K/R|X) and arginine-only cleavage (R|X) (to account for inaccessible lysine residues due to TMT blocking), with up to 2 missed cleavages permitted. Fixed modifications included carbamidomethylation of cysteine (+57.021 Da), while dynamic modifications comprised methionine oxidation (+15.995 Da), phosphorylation (+79.966 Da) on serine, tyrosine, and threonine residues (only applied during phosphorylation-related searches), and TMT 16/18-plex labeling (+304.207 Da) on lysine residues and peptide N-termini. Precursor and fragment mass tolerances were set to ±20 ppm and ±0.02 Da, respectively, with HCD fragmentation; identifications were filtered at 1% FDR (PSM, peptide, and protein levels) using Percolator, requiring ≥1 unique peptide per protein and ≥1 PSM per peptide.

### Bioinformatics and Data Analysis

#### Global Proteomics

##### Preprocessing and protein quantification

PSM-level TMT reporter ion intensities were exported from Proteome Discoverer 3.0 and read from replicate-specific spreadsheets (S1-S3). The peptide-reference channel (TMT 126) was excluded from all downstream analyses. Intensities were log2-transformed using a pseudocount (log2(abundance + 1)) and mapped to brain regions via the predefined TMT channel-region assignment table. PSMs lacking valid UniProt accessions were discarded. Protein-level quantification was obtained by two-stage median aggregation: first, PSM intensities were summarized to peptide-level medians for each unique combination of protein accession, annotated sequence, and sample; second, peptide-level values were summarized to protein-level medians for each accession-sample combination. The resulting protein × sample matrix was median-centered within each sample by subtracting the column-wise median, placing all samples on a common intensity scale prior to differential analysis.

To ensure reliable protein-level measurements, proteins were retained only if they had ≥2 finite quantified values in at least one brain region across replicates. This per-region reproducibility filter eliminates proteins present in a single sample but lacking cross-replicate support. Batch effects across the three biological replicates were removed using limma’s removeBatchEffect, with replicate set (S1–S3) as the batch factor and a region-only design matrix to preserve biologically meaningful regional signal. This batch-corrected matrix was used exclusively for visualization, unsupervised analyses, and the brain region heatmap viewer. All differential abundance analyses were performed on the non-batch-corrected, median-centered matrix with batch effects modeled explicitly as a linear covariate.

For the heatmap viewer, a region summary matrix was computed from the batch-corrected protein matrix by averaging across replicates within each region using a 20% trimmed mean, which downweights outlier values without discarding data. Proteins with insufficient regional coverage after aggregation were excluded, yielding a final set of 7,080 proteins with reliable spatial measurements across all ten regions.

##### Differential protein abundance analysis

Differential abundance testing was performed using the limma linear modeling framework on the median-centered protein matrix. Proteins quantified in ≥70% of samples were retained. A design matrix was constructed with region terms and a replicate batch covariate (S1-S3). For each of the ten brain regions, a contrast was defined as that region’s mean against the unweighted mean of all other nine regions. Contrasts were specified using makeContrasts and fitted using contrasts.fit, followed by empirical Bayes moderation (eBayes). Multiple testing correction was applied using the Benjamini–Hochberg (BH) method across all features within each contrast. Proteins with adjusted p-value < 0.05 and |log2FC| > 1 were considered significantly region-enriched. The top 20 most significantly enriched proteins per region were annotated on volcano plots.

##### Protein co-expression modules and cell-type composition

To identify coordinated spatial protein programs, protein profiles were summarized to region means (averaging across S1-S3 from the batch-corrected matrix), filtered to retain proteins with ≤30% missingness across regions, and z-scored per protein across regions. Mfuzz soft clustering was applied to the resulting matrix using the ExpressionSet framework; the optimal fuzzification parameter m was estimated using the mestimate function, and k = 8 clusters were selected based on elbow analysis of within-cluster sum of squares. Module membership scores (ranging 0-1) were retained for each protein, with module assignment based on the maximum membership value. Module-level activity scores were computed as the mean z-score of member proteins per region and visualized as profile plots and heatmaps. Functional enrichment of module protein sets was performed using Gene Ontology Biological Process analysis via fgsea, after mapping UniProt accessions to mouse gene symbols using org.Mm.eg.db. Cell-type composition across brain regions was assessed by computing the mean z-score of curated marker proteins across eight cell-type categories (Synaptic, Oligodendrocyte, Myelin, Microglia, Inhibitory, Excitatory, Endothelial, Astrocyte)^28–32^, which are included in the full protein matrix and whose cluster distributions reflect region-specific cell-type identity.

#### Phosphoproteomics

##### Preprocessing and quantification

PSM-level TMT reporter ion intensities were exported from Proteome Discoverer 3.0 and read from replicate-specific spreadsheets (S1-S3). Phosphosite-level quantitative tables from three independent replicates (S1-S3) were generated in a similar fashion. The reference channel (TMT 126) was excluded. Phosphosite identifiers were parsed to extract UniProt accessions and residue positions, and gene symbols were mapped using org.Mm.eg.db. A phosphosite × sample matrix was constructed. For differential analysis, phosphosites quantified in ≥70% of samples were retained; replicate effects were modeled explicitly via the batch covariate in the design matrix. For visualization, batch effects were removed using limma’s removeBatchEffect with a region-only design matrix. All differential testing used the non-batch-corrected matrix.

For the heatmap viewer, phosphosite region means were computed by pooling all three replicates before averaging, and sites quantified in ≥4 of 10 regions were retained. This union-based approach recovers sites detected in any replicate and expands coverage to 3,676 phosphosites (substantially more than the 2,456-2,552 identified per individual replicate), while excluding sites with insufficient regional representation.

##### Region-enrichment scores and protein-level correction

A direct region-enrichment score was computed as a continuous summary for KSEA and spatial clustering. For each phosphosite and each brain region, the enrichment score was defined as the difference between mean regional log2 abundance and the mean across all other regions, computed from the median-centered phosphosite matrix. Scores were set to NA when fewer than 2 finite values were available within the region or fewer than 5 across all other regions, ensuring reliable estimation. This approach yields finite scores for all sites with any quantification, enabling downstream analysis of the full phosphoproteome rather than the sparse subset passing the limma rank-sufficiency requirement.

Protein-level correction was performed to distinguish phosphorylation changes driven by altered protein abundance from those reflecting genuine site-specific kinase regulation. For each phosphosite with a matched parent protein in the global proteome matrix, a protein-corrected matrix was constructed by computing the element-wise difference between the median-centred log2 phosphosite abundance and the median-centred log2 abundance of the matched parent protein, aligned to the same sample order. NA values propagate where either the phosphosite or protein value is missing. The protein-corrected matrix was then subjected to the same limma differential analysis pipeline (design with region and batch terms, contrasts for each region vs. all others, eBayes, BH FDR correction). Phosphosites were classified into four regulatory categories based on significance in the uncorrected and protein-corrected analyses: (i) expression-dominant: significant in uncorrected analysis only (adj.P < 0.05, |log2FC| > 1), indicating phosphorylation change mostly accounted for by protein abundance; (ii) expression-masked: significant in protein-corrected analysis only, indicating regulation detected only after removing the protein-abundance component; (iii) regulation-dominant: significant in both analyses, indicating robust regulation beyond protein-level changes; (iv) not classified: significant in neither.

##### Kinase Substrate Enrichment Analysis (KSEA)

Kinase-substrate relationships were retrieved from the OmniPath enzyme-substrate database restricted to phosphorylation events in Mus musculus (all resources), and enzyme and substrate accessions were mapped to gene symbols using org.Mm.eg.db. For each brain region, the region-enrichment score vector served as the phosphosite-level activity measure. Where multiple phosphosites mapped to the same gene, the site with the maximum absolute enrichment score was retained as the representative value. A KSEA score was computed for each kinase as the scaled difference between the mean enrichment score of its detected substrates and the background mean across all quantified substrate genes, weighted by the square root of the number of detected substrates. Statistical significance was assessed by two-tailed z-test with BH FDR correction applied within each region independently. Kinases with fewer than two detected substrates in each region were excluded from testing. For heatmap visualization, kinases with zero variance across regions were removed; kinases untested in a given region were imputed to that kinase’s median score across quantified regions prior to hierarchical clustering (Ward’s D2 linkage on Euclidean distance) but displayed as grey in the final heatmap. FDR-significant kinase-region pairs (adj.P < 0.05) were marked with an asterisk overlay.

##### Spatial phosphosite modules (Mfuzz soft clustering)

To identify groups of phosphosites with coordinated spatial regulation across brain regions, Mfuzz soft clustering was applied to region-enrichment score profiles. Sites were required to have finite scores in ≥6 of 10 regions; remaining missing values were imputed by row median. Profiles were z-scored across regions (row-wise scaling), and sites with any non-finite z-scores after scaling were excluded. The top 50% most variable sites (by row variance of z-scored profiles) were retained for clustering, reducing noise from low-variance sites. The fuzzification parameter m was estimated using the mestimate function applied to an ExpressionSet object. The number of clusters k was selected by elbow analysis of within-cluster sum of squares (k-means, 25 restarts, 500 iterations; evaluated for k = 4 to min(16, floor(n/10))). Mfuzz clustering was run with set.seed(1) for reproducibility, yielding six fuzzy phospho-modules (PM1-PM6) with continuous membership scores for each site in each module. Module assignment was based on maximum membership value. Module membership scores and site-level assignments were exported for downstream pathway enrichment and Sankey visualization.

##### Kinase-Phosphosite-Pathway-Brain Region Sankey integration

A four-layer Sankey diagram was constructed to integrate kinase module membership, phospho-module membership, enriched Reactome pathways, and brain region distributions into a unified view of spatial signaling architecture. Kinase modules (K-Mod 1-4) derived from Mfuzz clustering of KSEA scores in Section 1 were annotated with KinBase structural and evolutionary group classifications using regex-based pattern matching against kinase gene symbols, with the top two represented groups used as module labels.

Edges between kinase modules and phospho-modules were computed by mapping each kinase module’s member kinases through the OmniPath kinase-substrate database to their detected substrate phosphosites, then counting the number of distinct phosphosites connecting each kinase module to each phospho-module; edges with fewer than two distinct phosphosites were excluded. Reactome pathway annotations for Mus musculus were retrieved from MSigDB (C2:CP:REACTOME) via msigdbr, and phosphosites were mapped to pathways through their parent gene symbols. For each phospho-module, the top four pathways by number of distinct member phosphosites were retained. Edges from phospho-modules to brain regions were weighted by the mean z-score of module member sites in each region, rectified at zero and normalized to proportions summing to one per module. The composite flow value for each four-way combination was computed as the product of the normalized kinase-module-to-phospho-module weight, the normalized phospho-module-to-pathway weight, and the region proportion. To reduce visual clutter while preserving representation of all brain regions, flows were filtered by retaining only those above the 60th percentile within each region independently.

## Resource availability

### Lead contact

Further information and requests for resources should be directed to and will be fulfilled by the lead contact, Hui Zhang.

### Materials availability

This manuscript contains no unique reagents or resources. All reagents, tissue materials, and instruments are available commercially.

### Data and code availability

The mass spectrometry raw data have been deposited to the ProteomeXchange Consortium via the PRIDE partner repository with the dataset identifier PXD077317.

Any additional information required to reanalyze the data reported in this paper is available from the lead contact upon request.

## Acknowledgement

This work was supported by National Institutes of Health, National Cancer Institute, Early Detection Research Network (EDRN, U2C CA271895), Pancreatic Cancer Detection Consortium (PCDC, U01 CA274514), and Clinical Proteomic Tumor Analysis Consortium (CPTAC, U24 CA271079). We thank Dr. Heng Zhu (Johns Hopkins University, Department of Pharmacology and Molecular Sciences) for generously providing access to the GenePix 4000B Microarray Scanner (Molecular Devices).

## Author contributions

YX, JL, and HZ conceived and designed the study. YX, JL, HCP, YL, CYL, XL and HZ contributed to the design and modification of the 3D printing platform. YX optimized the printing platform, developed and optimized the SPOTTER proteomics workflow. YX performed experiments, conducted data analysis, and interpreted the results. TML provided guidance on data analysis. YX wrote the original draft of the manuscript. All authors reviewed, edited, and approved the final manuscript.

---

**Figure S1.**
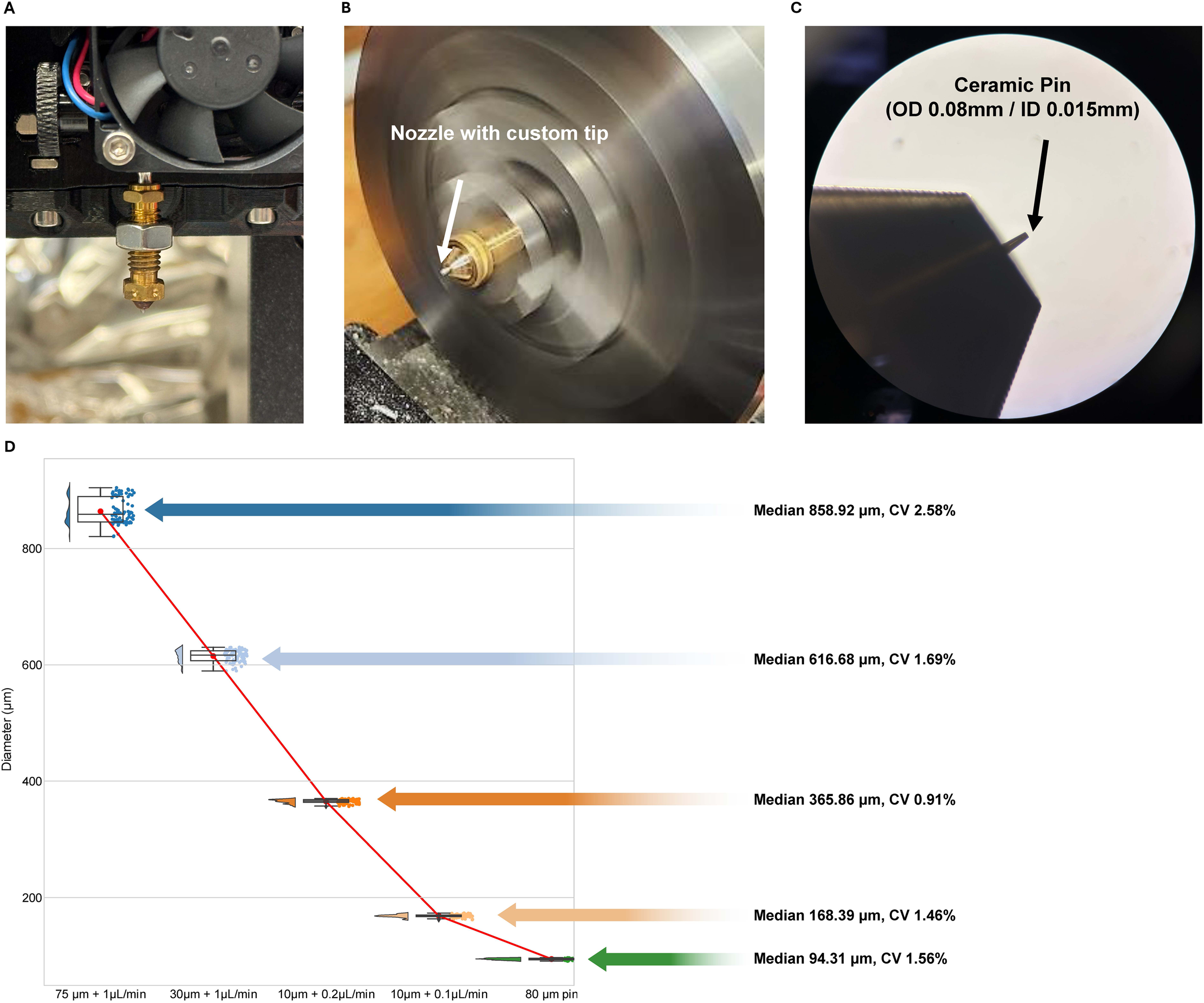
SPOTTER print head assembly and printing performance evaluation. A) Photograph of the SPOTTER print head assembly. B) Microscopic image of the print head nozzle coupled to a fused silica nano-flow liquid chromatography column. C) Microscopic images of the pin-based printhead nozzle. D) Printing performance evaluation across three nozzle inner diameters (75, 30, and 10 μm) and corresponding flow rates (1, 0.2, and 0.1 μL/min), together with an additional 80 μm-i.d. pin-based printing, assessed by depositing a 10 × 10 grid pattern of a fluorescent solution (1% rhodamine B in 25% v/v acetonitrile). The jitter plot shows the distribution of measured dot diameters across all conditions, with median diameter and coefficient of variation (CV) annotated for each.

**Figure S2.**
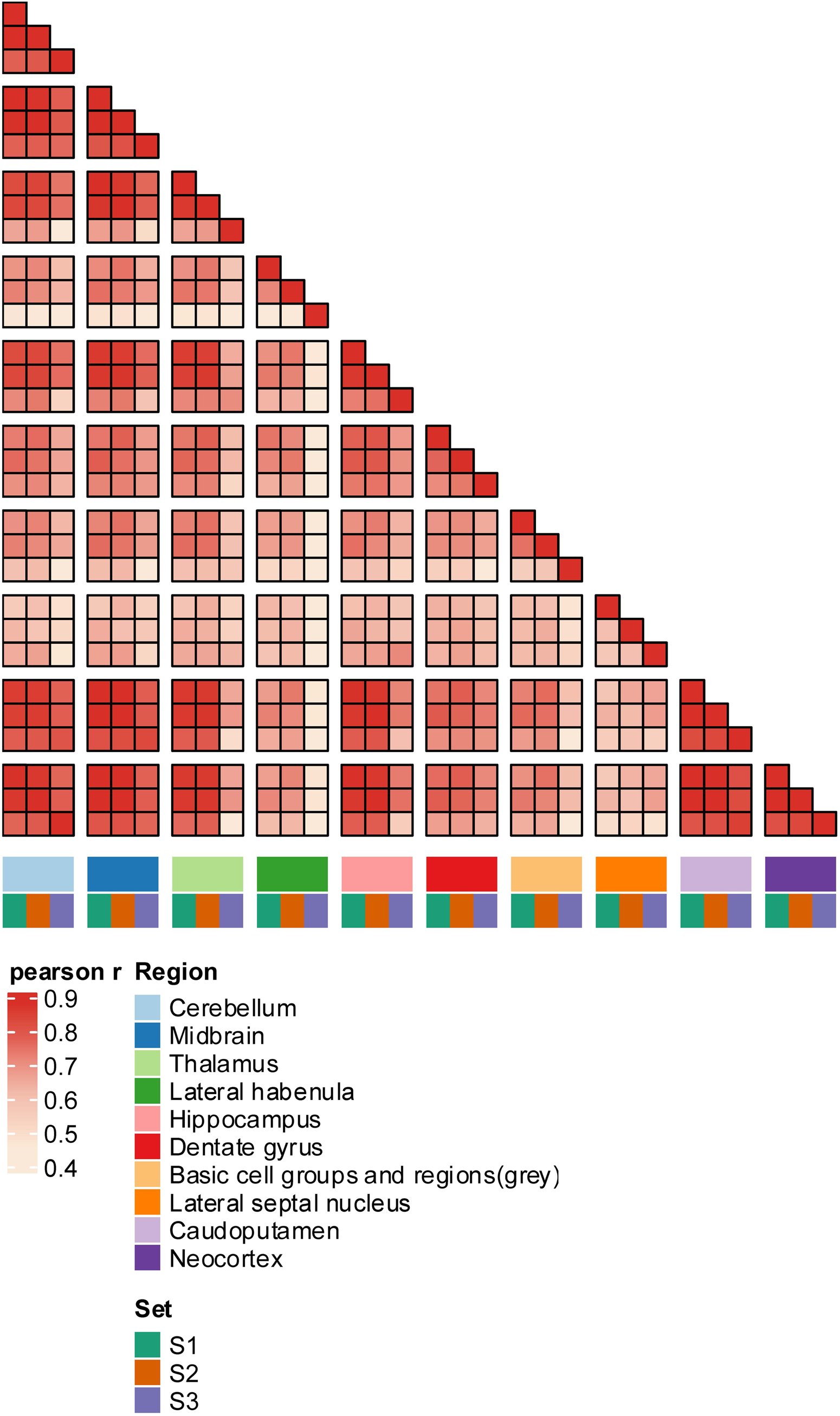
Reproducibility of spatial proteomics data. Pairwise Pearson correlation matrix of protein abundance values across the three replicates, colored by brain region and replicate set, demonstrating high inter-replicate reproducibility.

**Figure S3.**
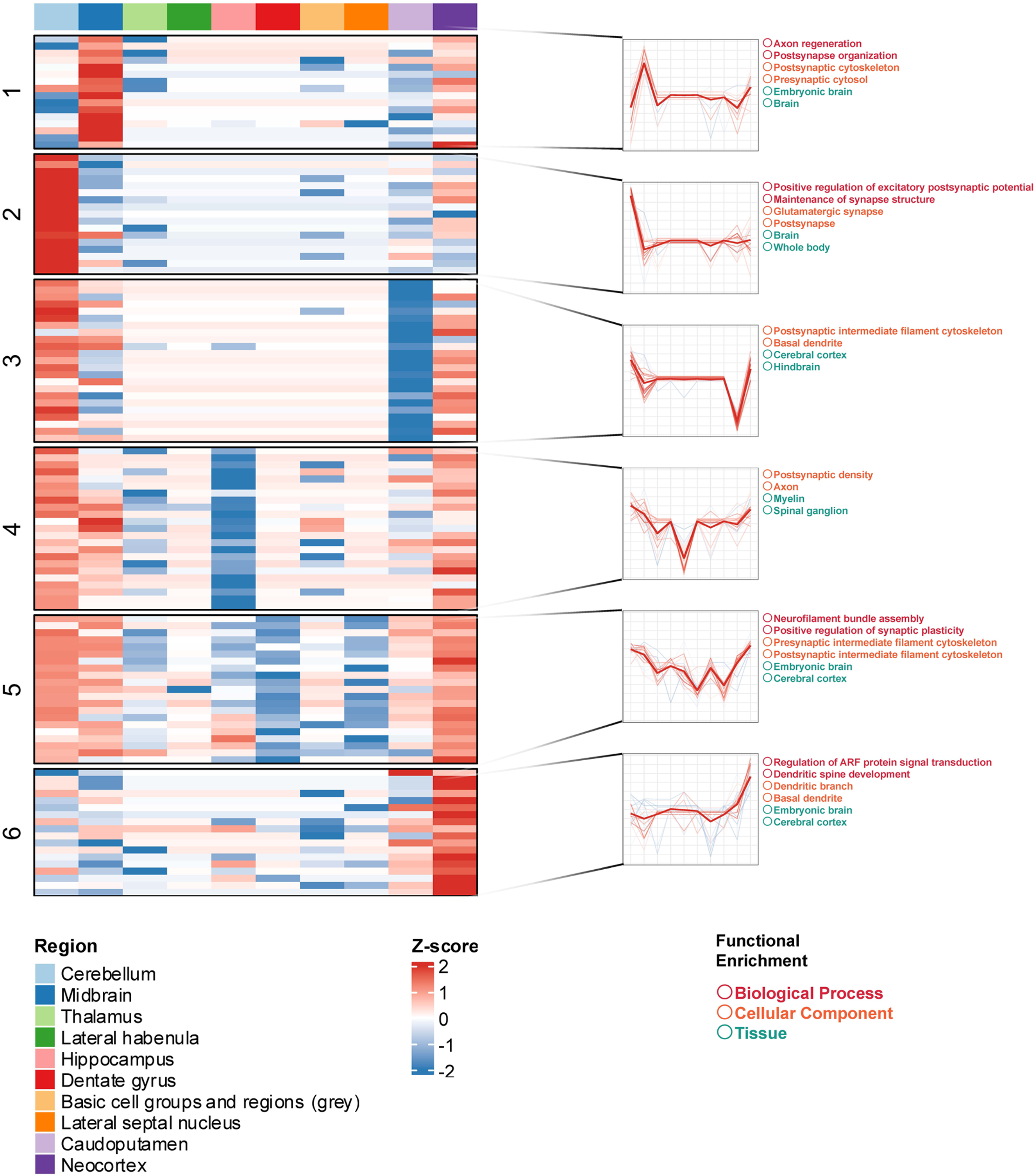
Soft clustering of phosphoproteomics data across ten brain regions. Soft clustering of phosphoproteomics data across ten brain regions using Mfuzz yields six fuzzy clusters, displayed as Z-score heatmaps. Cluster membership profiles are shown as line plots (center), and functional enrichment terms from biological pathway, cellular component, KEGG pathway, and tissue ontologies are annotated for each cluster (right).

**Figure S4.**
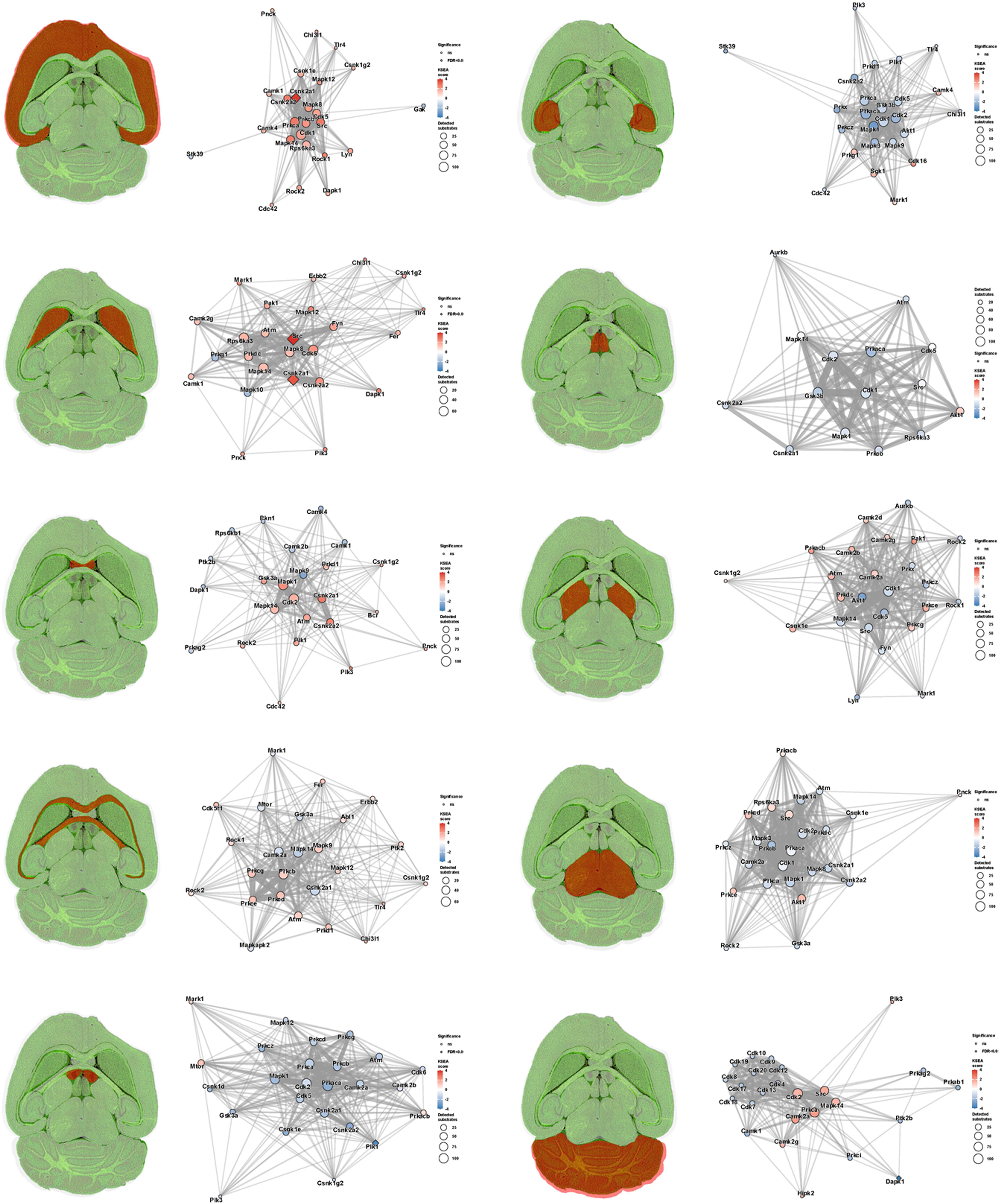
Region-wise kinase activity networks across all ten brain regions. For each brain region, a brain atlas overlay (left) highlights the region of interest (red) relative to all other regions (green), paired with a kinase co-regulation network (right) showing the top 25 kinases ranked by absolute KSEA score. Nodes are colored by KSEA activity score (red = activated, blue = inhibited) and sized by the number of detected substrates. Edges connect kinase pairs that share one or more detected substrates, with edge width proportional to the number of shared substrates.

## Notes

### Competing Interest Statement

The authors have declared no competing interest.

